# High-confidence calling of normal epithelial cells allows identification of a novel stem-like cell state in the colorectal cancer microenvironment

**DOI:** 10.1101/2024.02.23.581690

**Authors:** Tzu-Ting Wei, Eric Blanc, Stefan Peidli, Philip Bischoff, Alexandra Trinks, David Horst, Christine Sers, Nils Blüthgen, Dieter Beule, Markus Morkel, Benedikt Obermayer

## Abstract

Single-cell analyses can be confounded by assigning unrelated groups of cells to common developmental trajectories. For instance, cancer cells and admixed normal epithelial cells could potentially adopt similar cell states thus complicating analyses of their developmental potential. Here, we develop and benchmark CCISM (for Cancer Cell Identification using Somatic Mutations) to exploit genomic single nucleotide variants for the disambiguation of cancer cells from genomically normal non-cancer epithelial cells in single-cell data. In colorectal cancer datasets, we find that our method and others based on gene expression or allelic imbalances identify overlapping sets of cancer versus normal epithelial cells, depending on molecular characteristics of individual cancers. Further, we define consensus cell identities of normal and cancer epithelial cells with higher transcriptome cluster homogeneity than those derived using existing tools. Using the consensus identities, we identify significant shifts of cell state distributions in genomically normal epithelial cells developing in the cancer microenvironment, with immature states increased at the expense of terminal differentiation throughout the colon, and a novel stem-like cell state arising in the left colon. Trajectory analyses show that the new cell state extends the pseudo-time range of normal colon stem-like cells in a cancer context. We identify cancer-associated fibroblasts as sources of WNT and BMP ligands potentially contributing to increased plasticity of stem cells in the cancer microenvironment. Our analyses advocate careful interpretation of cell heterogeneity and plasticity in the cancer context and the consideration of genomic information in addition to gene expression data when possible.

**Novelty and Impact:** Single-cell analyses have become standard to assess cell heterogeneity and developmental hierarchies in cancer tissues. However, these datasets are complex and contain cancer and non-cancer lineage cells. Here, we develop and systematically benchmark tools to distinguish between cancer and non-cancer single-cell transcriptomes, based on gene expression or different levels of genomic information. We provide strategies to combine results of different tools into consensus calls tailored to the biology and genetic characteristics of the individual cancer.

## Introduction

Cancer cells mix and interact with their microenvironment ^1,2^. In colorectal carcinoma (CRC) and in other epithelial cancers, transformed cells intermingle with non-cancer epithelial cells in areas known as the invasive front (IF)^3,4^. Furthermore, normal tissues adjacent to tumours are re-shaped beyond the cancer’s boundary, influenced by local immune responses and inflammation ^5^, paracrine signals ^6^, and genetic aberrations preceding malignant transformation ^7^, as has been shown by multiplexed tissue imaging ^8^, single-cell ^9^ and bulk transcriptomics ^10^. This gradual change in cell composition from normal to cancer poses challenges for single-cell transcriptomics, as it is not immediately apparent from the transcriptome whether certain cells arise from malignant or normal lineages.

In CRC, single-cell transcriptome analyses revealed two overarching intrinsic consensus molecular subtypes (iCMS), termed iCMS2 and iCMS3^11^. These transcriptome subtypes are linked to patient characteristics such as localization of cancer, and to molecular features such as microsatellite stability, mutational burden, the extent of copy number aberrations, and patterns of driver mutations ^12–15^. That means, left-sided tumours frequently arise due to the loss of the tumour suppressor gene APC and additionally harbour mutations in KRAS, SMAD4, and TP53; these mutational patterns lead to WNT and MYC signalling pathway activation. Furthermore, CRCs in this context are most frequently microsatellite-stable (MSS), display extensive copy number aberrations and gene expression patterns characteristic of intrinsic molecular subtype iCMS2. In contrast, CRCs progressing via serrated precursors are found mainly in the right colon, carry mutations in KRAS or BRAF, display activation of the TGF-beta signalling pathway, can be microsatellite-instable (MSI) or MSS, have a higher mutational burden but fewer copy-number changes, and show gene expression patterns of metaplasia and intrinsic molecular subtype iCMS3. We expect that the different cancer cell characteristics could also lead to a variable accuracy of cell type calling in single-cell analysis.

Numerous studies have conducted single-cell level analyses of CRC ^16–18^. These investigations were either performed under the assumption that all epithelial cells derived from the cancer tissue samples are bona fide cancer cells, or they have relied solely on transcriptome-derived characteristics to differentiate between cancer and normal epithelial cells. Broadly applicable and robust methods to confidently distinguish genomically normal epithelial cells from genomically aberrant cancer cells remain elusive, especially for datasets derived from regions where both types of cells coexist, such as at the IF.

Here, we use different computational tools to disambiguate cancer and non-cancer epithelial cells in single-cell transcriptome data of ten CRC patients across a range of clinical and molecular characteristics, using additional information derived from associated whole-genome sequencing data. Analysis of consensus sets of cancer and normal cells shows that genomically normal epithelial cells adjacent to the cancer can adopt cell states that are unlike those of epithelial cell populations in normal tissue. Developmental trajectories of non-cancer epithelium were altered in the cancer neighbourhood, as stem-like and immature differentiation states were overrepresented among genomically normal cells in cancer tissue samples. We identify multiple new paracrine interactions potentially modulating normal cell development in the tumour microenvironment, including cancer-specific fibroblasts as a source of the key stemness factor WNT.

## Results

### Transcriptome information is insufficient for cancer cell calling in CRC

To reliably distinguish cancer from normal cells in single-cell RNAseq data, we complemented single-cell data of ten treatment-naive CRC patients of a previous study ^17^ with whole-genome sequencing data of cancer and normal samples. Clinical and pathology assessment of the cohort shows a broad distribution along the longitudinal axis of the colon, and driver mutations in APC, BRAF, P53, beta-Catenin, and KRAS in subsets of the cancers (Fig. 1A). Using updated bioinformatic pipelines, 73 294 cells passed quality controls after ambient RNA and doublet removal. Of these, 43 110 transcriptomes were from cancer tissue and 30 184 from normal tissue samples adjacent to tumour. 39 168 cells were annotated as epithelial, 31 663 as immune cells and 2 463 as of stromal cell origin (Fig. 1B).

**Figure 1.**
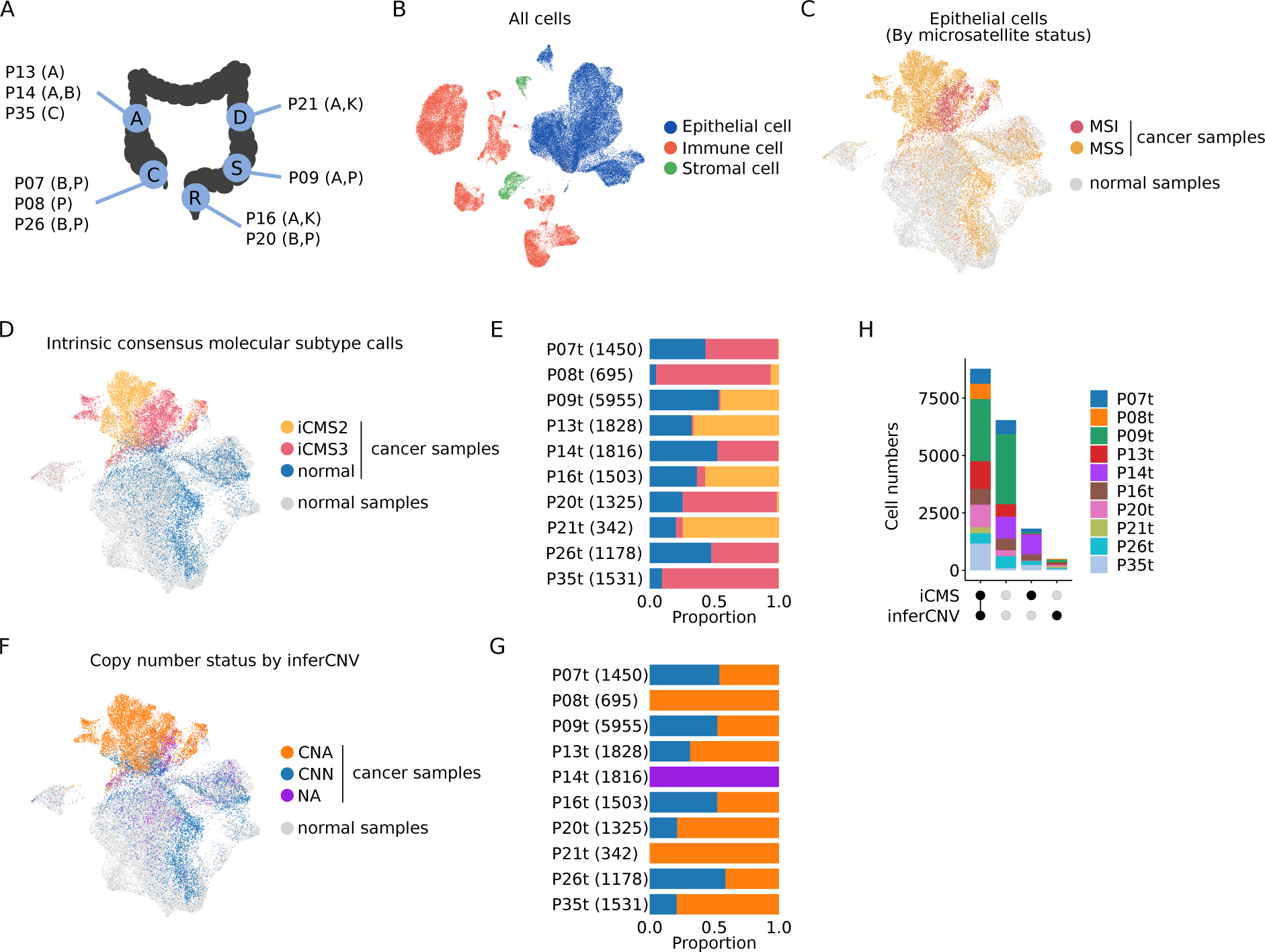
Cancer cell calling based on transcriptome information. **A** Anatomical locations and mutational patterns of the samples. C: cecum, A: ascending colon, D: descending colon, S: sigmoid, and R: rectum. Mutations (in brackets) A: *APC*, B: *BRAF*, C: *CTNNB1*, K: *KRAS*, P: *TP53*. **B** UMAP of all 73 294 cells, coloured by three major cell type compartments: epithelial (blue), immune (orange), and stromal cells (green). **C, D, F** UMAPs of epithelial cells only. (C) colour code by the sample origin and the microsatellite status. Cancer sample (MSI), red; cancer sample (MSS), yellow; normal sample, grey. (D) colour code for cancer sample cells by iCMS assignment; iCMS2 (yellow), iCMS3 (pink), or normal (blue), normal samples (not scored, grey). (F) colour code of cancer sample cells by inferCNV. Copy number status aberrant (CNA; orange), normal (CNN; blue), or not applicable (NA; purple) when the clones in the sample are not differentiable, normal samples (not scored, grey). **E, G** stacked bar plots summarising iCMS and inferCNV information, respectively, by cancer sample. **H** Quantification of the agreement between iCMS and inferCNV calls as an upset plot, colour-coded by patient, as indicated.

We first sought to distinguish cancer from normal epithelial cells in the cancer samples using transcriptome information. In a UMAP representation of all epithelial cell transcriptomes, a fraction of the 17 623 transcriptomes derived from cancer samples partially clustered on a separate “community” while another fraction interspersed with the normal tissue-derived epithelial cells (Fig 1C). We used probabilistic label transfer from published gene expression data^11^ to assign cancer sample epithelial cells to the cancerous iCMS2 or iCMS3 epithelial cell states, or a normal cell state (Fig. 1D, E). In total, 10 589 cells were classified as iCMS2 or iCMS3, and therefore were assigned as cancer cells by this method. Cancer cells from P09, P13, P16 and P21 received predominantly called as iCMS2, whereas P07, P08, P14, P20, P26 and P35 were mostly iCMS3, in line with previous analyses showing that MSI cancers are usually iCM3. However, our analysis also showed that cells displaying transcriptome features of iCMS2 and iCMS3 can be present side-by-side in the same cancers (Fig. 1E). Almost all the cells receiving iCMS2/iCMS3 calls were located on the cancer cell community of the UMAP in contrast to the 7 034 cancer tissue-derived epithelial cells receiving the “normal” label that were mostly scattered among cells derived from normal tissue samples.

We also inferred cancer cell identity by expression-derived copy number calls, using inferCNV ^19^ (Fig. 1F,G). Using hierarchical clustering based on copy number-driven genome-averaged expression patterns (Fig. S1), we assigned cell clusters as cancer when their averaged expression pattern deviated more than three standard deviations from epithelial cells in the normal tissue samples. This method did not yield results for the MSS cancer P14, which did not exhibit detectable alterations in the averaged expression patterns. For the remaining cancer samples, inferCNV identified a total of 10 509 abnormal transcriptomes, whereas 7 114 cells were assigned as normal epithelial cell states.

Taken together, the transcriptome-based analyses showed a large overlap for calling cancer versus genomically-normal cells (Fig. 1H). However, 1 441 cells received conflicting calls and cells from P14 could not be properly assigned. Thus, these methods are not suitable to generally define genomically-normal versus cancer epithelial cells in CRC samples with high accuracy.

### Exploiting cancer specific SNV information for cancer cell calling with CCISM

Given that transcriptome analyses can potentially be confounded by expression similarities between cancer and normal epithelial cell states, we hypothesized that independently derived somatic variants that are observed in single-cell sequencing reads constitute the most unambiguous evidence that a cell originated from a cancer lineage. We therefore utilised cancer-specific somatic variants derived from bulk whole-genome sequencing data of matched samples to interrogate the associated single-cell transcriptomes.

Comparison of normal and cancer genomes yielded 2-12 cancer-specific somatic single nucleotide variants (SNVs) per million bases of genome sequence (MB) in most CRCs, except for the MSI CRCs P26 and P35 which had up to 50 SNVs/MB (Fig. 2A). The mean number of expressed SNVs per cell in the single-cell transcriptomes correlated with the SNV frequency in the whole-genome sequencing data and was for many CRCs less than 10 SNVs per cell, but up to 60 SNVs/cell for the MSI CRC P35.

**Figure 2.**
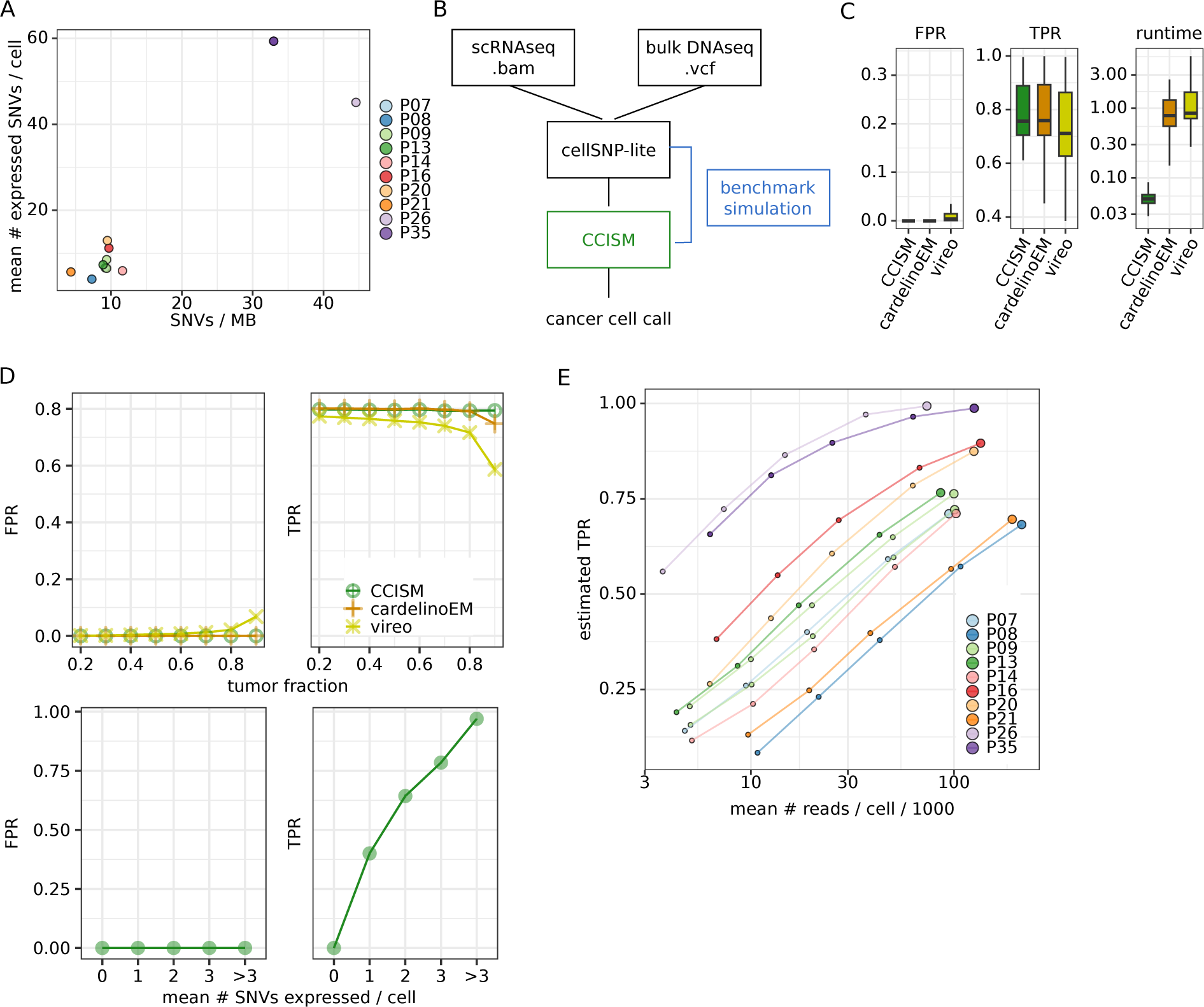
CCISM identifies cancer cells with somatic single nucleotide variants. **A** Scatterplot of the number of SNVs in whole genome sequencing data and the average number of expressed SNVs per cell in single-cell RNA sequencing data coloured by patient. **B** CCISM’s workflow diagram from input data (scRNAseq and bulk DNAseq data), allele count calculation by cellSNP-lite to CCISM modelling. Benchmark simulations can be generated from input counts (blue). **C** Boxplots of tool performances in simulation data regarding runtime in seconds (right), false positive rate (FPR, middle), and true positive rate (TPR, left) between CCISM (green), cardelinoEM (orange), and vireo (pear). **D** Line plots comparing model performances (CCISM, green circle; cardelinoEM, orange cross; vireo, pear star) as function of tumour fraction (upper) and mean number of expressed SNVs per cell (lower). **E** Line plot of CCISM’s performance (TPR) in single-cell transcriptomes subsampled to five different mean numbers of reads per cell, colour-coded by patient.

To make use of SNV patterns for the classification of single-cell data, we developed CCISM (for Cancer Cell Identification by Somatic Mutations). Input data are the UMI-collapsed read counts for reference and alternative allele observed per cell and variant, which are obtained from the single-cell sequencing reads as well as a list of high-quality somatic variants derived from bulk whole-genome or whole-exome data. Based on this input, CCISM computes for each cell a posterior cancer cell assignment by expectation maximization. Importantly, these are cell-specific values and not derived from clustering. At the same time, benchmark simulations can be used to estimate expected sensitivity and specificity values for the dataset at hand (Fig. 2B).

We first used simulations based on the total allele count matrices from our single-cell RNAseq datasets to benchmark CCISM against cardelinoEM ^20^ and vireo ^21^. Compared to these existing tools with related functionality, CCISM has similar specificity but superior computational efficiency (Fig. 2C). We also obtained better sensitivity especially at high tumour content, mainly because we employed a fixed parameter for the probability to observe variant alleles in normal cells instead of estimates. Note that sensitivity depends on the number of expressed SNVs per cell and reaches optimal values at three or more expressed SNVs per cell (Fig. 2D). Across the datasets used to initiate the simulations, we found sensitivity strongly associated with mutational burden and therefore highly correlated to the average number of expressed SNVs per cell (Fig. S2A). A subsampling analysis revealed that most datasets were not saturated for SNV coverage despite being sequenced to depths of more than 90 000 autosomal reads per cell on average (Fig. 2E).

### CCISM and Numbat can be used cooperatively to define consensus normal and cancer cell lineage populations

We applied CCISM to our CRC single-cell RNA dataset resulting in 9 738 cancer cell calls (Fig. 3A). The predicted cancer cells show a widely overlapping localization with cells previously classified using expression-based copy-number variation inference with inferCNV or iCMS2/iCMS3 gene expression (Fig. 1D, F). However, CCISM generated more cancer cell calls in UMAP neighbourhoods identified as normal by iCMS or inferCNV (Fig. 3A, see rectangular insets), suggesting that the use of cancer-specific variant information retrieves cells of cancer lineages that are transcriptomically less divergent from genomically normal epithelial cells.

**Figure 3.**
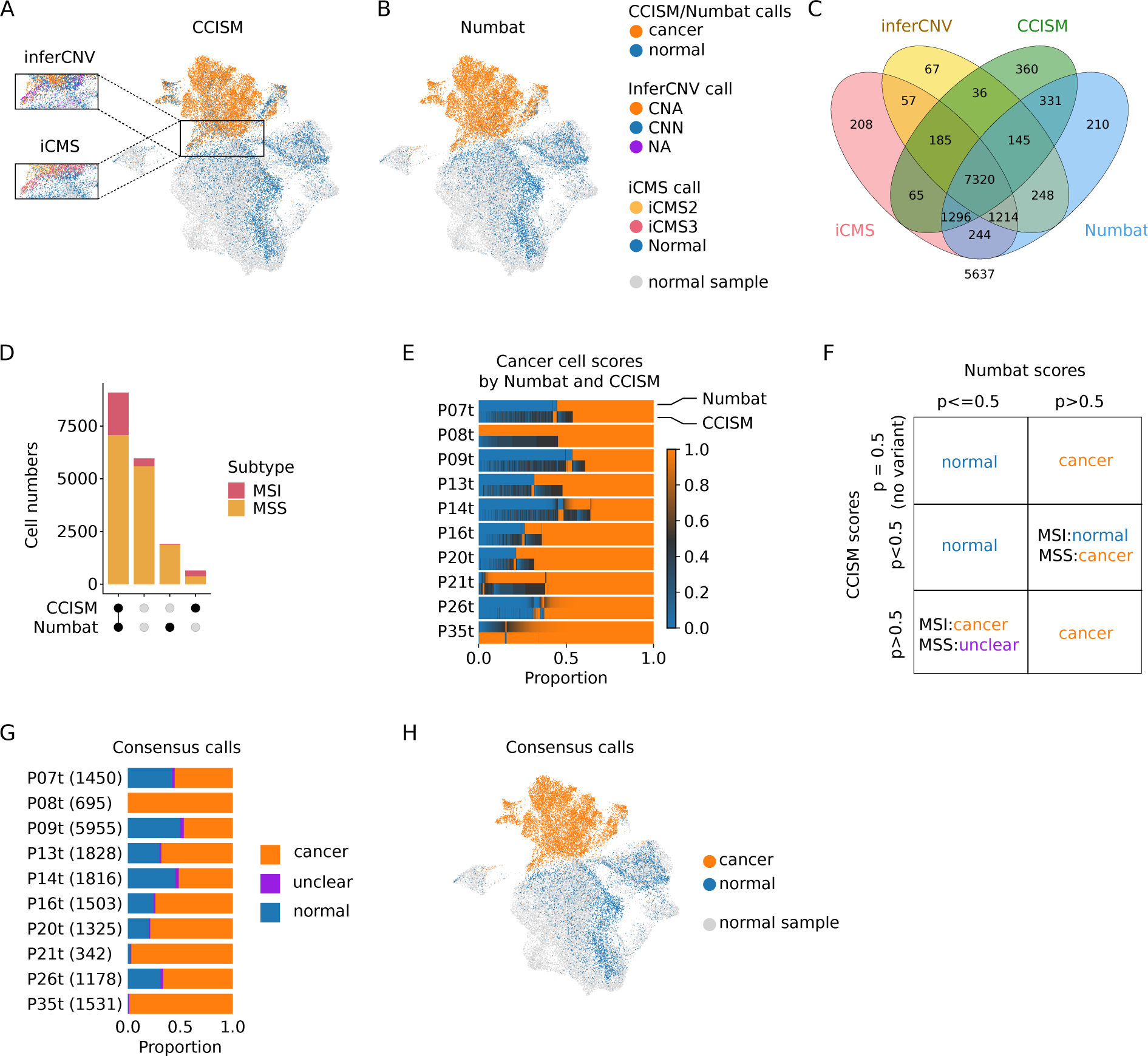
Cancer cell calling based on genomic information. **A, B** UMAPs of epithelial cells. **A** Colour-code by CCISM calls (cancer cell, orange; normal cell, blue). Insets given for inferCNV and iCMS calls. Cells from normal samples are given in gray. **B** Colour-code by Numbat call (cancer cell, orange; normal cell, blue). Cells from normal samples are given in gray. **C** Venn diagram of the intersections of cancer cell calls from iCMS (pink), inferCNV (yellow), CCISM (green), and Numbat (blue). **D** Upset plot of the intersections of cancer cell calls from CCISM and Numbat coloured by microsatellite status of the sample (MSI, red; MSS, yellow). **E** Heatmaps of the cancer cell scores (0.0, blue; 0.5, dark grey; 1.0, orange) from Numbat (upper) and CCISM (lower) across cancer samples. **F** Decision matrix for consensus cancer cell calls, based on CCISM, Numbat and microsatellite status. **G** Stacked barplot of the consensus derived from CCISM and Numbat (cancer cell, orange; normal cell, blue; undefined, purple). **H** UMAP of the consensus calls, colour code as in G, excluding cells with an “unclear” call.

For comparison, we employed Numbat ^22^, a recently developed tool using allele frequency shifts of common germline variants to facilitate cancer cell calling via detection of copy number changes. In our single-cell dataset, Numbat identified 11 008 cells as of cancerous origin, again showing an incomplete overlap with cancer cells identified by the other methods (Fig. 3B, Fig. 3C).

Initially, 2 562 cells received conflicting assignments by CCISM and Numbat (Fig. 3D). Therefore, we studied strengths and weaknesses of both tools, considering individual tumour characteristics (Fig. 3D-E, Fig. S2B, S2C, S2D). On the one hand, we found that in MSS CRCs, most cells with a conflicting assignment were earmarked as cancer cells by Numbat; however, these cells did not receive a high-confidence cancer cell score by CCISM, as they contained only a median of one SNV, with 707 cells expressing no SNV at all. On the other hand, cells with conflicting assignment in MSI CRC samples mostly (272/314) received a high-confidence cancer cell score by CCISM, and these contained a median of 16 SNVs, while cancer cell scores computed by Numbat were generally low. Therefore, we developed a set of rules to arrive at a cancer cell consensus based on genomic information (Fig. 3F): epithelial cells of cancer samples receiving high scores (>0.5) by Numbat were assigned as cancer cells, except for cells of MSI cancers that were assigned as normal by CCISM (<0.5), which then received a normal call. Epithelial cells of cancer samples receiving high scores by CCISM (>0.5) were also assigned cancer cells, except when this call of MSS CRC cells conflicted with a low score by Numbat (<0.5), in which case the cell was called “unclear”. Using these consensus call rules, we were able to assign 11 238 cells as cancer cells (Fig. 3G, H). 570 of these were not recognized as cancer cells by iCMS transcriptional signatures or by inferCNV. 5 969 cells were assigned as derived from normal epithelial lineages, using SNV or haplotype information (Fig. S2C). A remaining set of only 416 cells was assigned as “unclear” and removed from further analysis, as they contained no reliable SNV or haplotype information.

### Consensus cell identity leads to higher homogeneity of transcriptome clustering and enables phenotypic comparison

The final cell assignment to cancer or normal lineages resulted in a substantial separation of the populations when visualized on the UMAP (Fig. 3H). We quantified distributions of consensus call cancer versus normal calls in the corresponding louvain cluster structure (Fig. 4A, Fig. 4B). We found that normal and cancer cell communities were best separated when using the consensus call, compared to relying on the different methods that use transcriptome or genomic information individually (Fig. 4C, Fig. S3A-D). Using the consensus annotation, cancer cells were distributed in a highly patient-specific manner, but genomically normal epithelial cells intermingled as well as epithelial cells derived from normal tissue samples (Fig. S3E). While the consensus call requires additional genomic data, the correspondence to the louvain cluster structure also implies that transcriptomes alone may contain sufficient information for the disambiguation of cancer and normal lineage epithelial cells, at least in our CRC single-cell data set.

**Figure 4.**
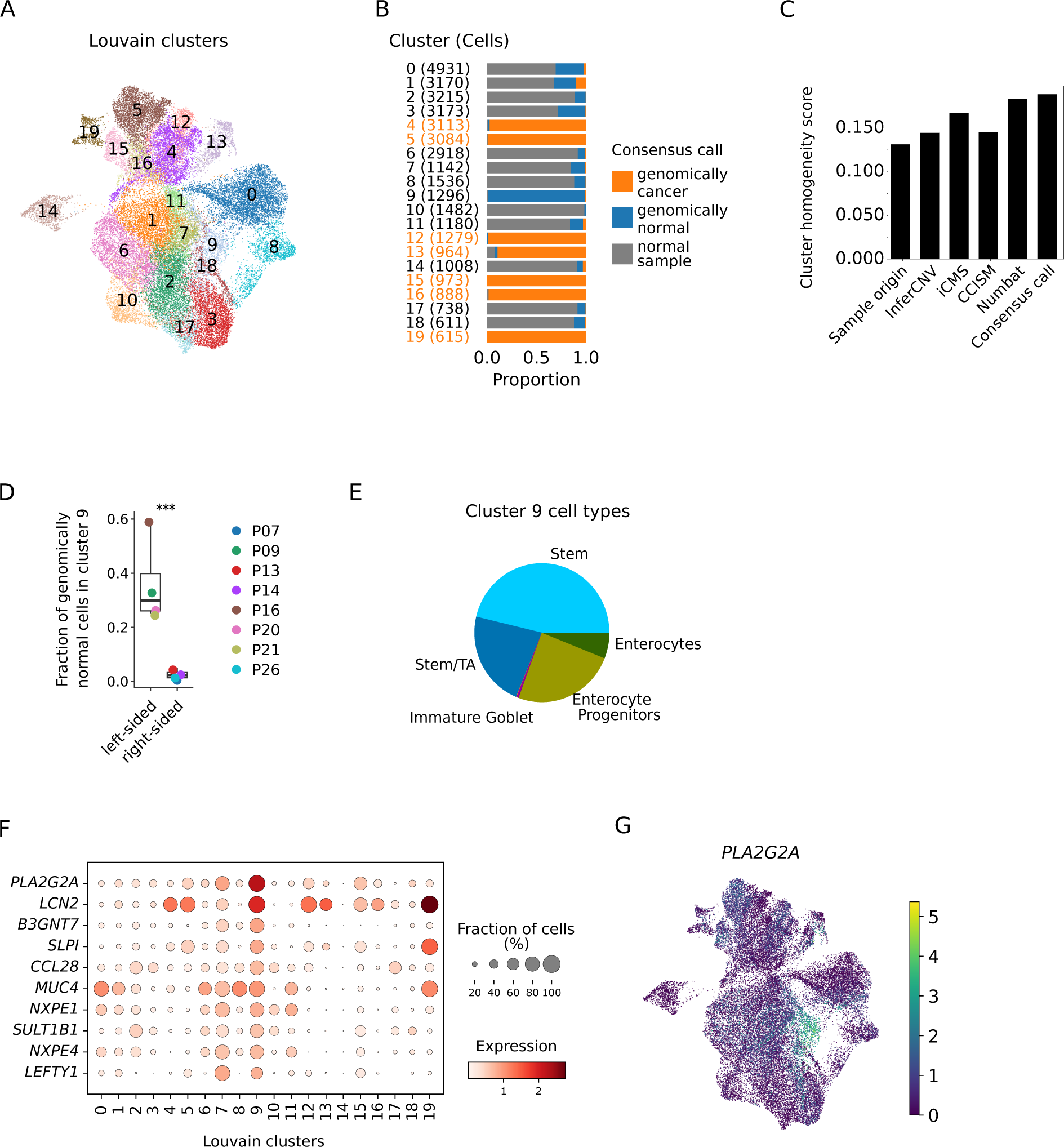
Consensus calls identify a cluster of genomically normal cells unique to left-sided cancer samples. **A** UMAP of epithelial cells, coloured by louvain clustering. **B** Stacked bar plot of consensus calls across 20 louvain clusters (cancer sample and genomically cancer, orange; cancer sample and genomically normal, blue, normal sample, grey). **C** Bar plot of cluster homogeneity scores for cancer cell calls by different methods as indicated. **D** Relative fractions of genomically normal cells in cluster 9, by cancer location (see Fig. 1A). p-value from mixed-effects binomial model, *** p < 0.001. **E** Pie chart of the epithelial cell types in louvain cluster 9, as indicated. Colour code: Enterocyte (dark green), Enterocyte progenitor (light green), Immature Goblet (light purple), Stem/TA (dark blue), and Stem (light blue). **F** Dot plot of top 10 marker genes for louvain cluster 9. Colour of dot represents the mean normalised expression of the gene, and the size of the dot shows the fraction cells expressing the gene. **G** UMAP coloured by *PLA2G2A* expression, which is the top gene marker specific to louvain cluster 9.

We found that the genomically normal epithelial cells from cancer samples showed distinct cluster distributions when compared to the normal tissue epithelial cells (Fig. 4B). In particular, louvain cluster 9 was almost exclusively composed of genomically-normal epithelial cells of cancer samples, and these were derived predominantly from tissue samples of patients P09, P16 and P20, and P21 with a left-sided (sigmoid colon and rectum) origin (Fig. 1A, Fig 4D). We further explored the identities of epithelial cells using label transfer ^17^. Cluster 9 contained mainly stem cells, transiently-amplifying cells or enterocyte precursors (Fig. 4E), and their assignment to a distinct louvain cluster suggested that these cells adopted a cell state that was induced by the cancer microenvironment and therefore not found in normal colon. When we analysed cluster 9-specific expression patterns, the most strongly defining gene for cluster 9 epithelial cells was *PLA2G2A*, encoding a secreted phospho-lipase (Fig. 4F, G).

Mapping of well-established colon and CRC cell-type signatures (Table S1) onto the epithelial single-cell transcriptomes derived from cancer and normal samples unveiled further differences in differentiation programs in the cancer’s vicinity, as Goblet cell transcriptomes derived from cancer samples were enriched for a Paneth cell signature, indicating that the cancer microenvironment perturbs secretory lineage fate decisions (Fig. S4A). Indeed, the occurrence of metaplastic Paneth cells has been widely documented in inflammation and also in cancer of the colon ^23,24^.

### The CRC microenvironment modulates epithelial cell states and developmental trajectories

We next assessed cell type frequencies among the genomically normal epithelial cells from cancer samples and compared them to normal tissue sample epithelium, excluding patients P08, P21, P26 and P35 which either had no matched normal sample or very few genomically normal cells (Fig. 5A). We found that the cancer-adjacent epithelial cells were significantly enriched for stem cells, immature goblet cells, and enterocyte progenitors, while they contained lower proportions of terminally differentiated cell types, such as differentiated enterocytes, goblet cells and tuft cells (Fig. S5B).

**Figure 5.**
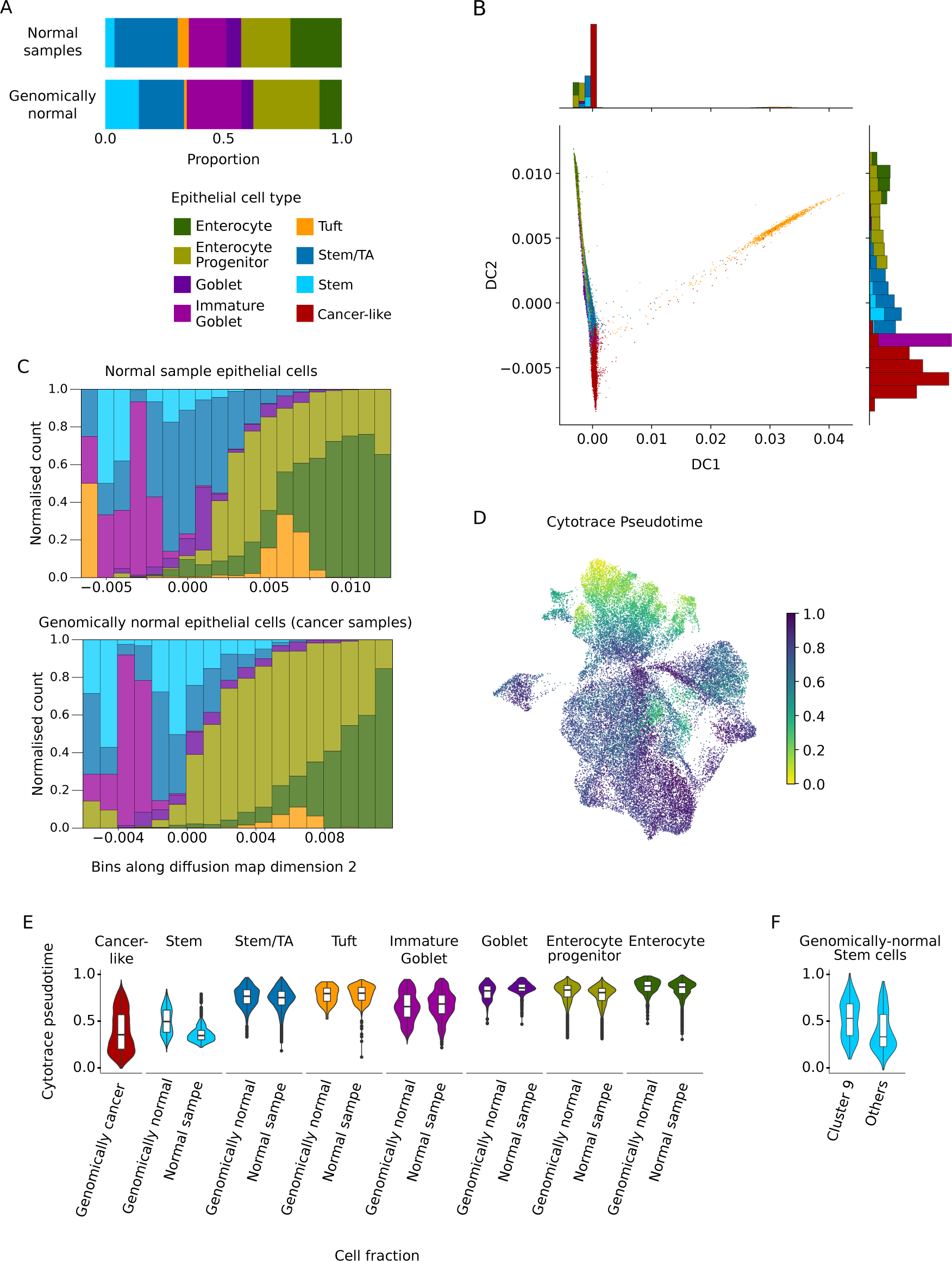
Cell states and developmental trajectories are altered in genomically normal cells of cancer samples compared to normal colon epithelium. **A** Stacked bar plots of epithelial cell types in normal samples (upper) and genomically normal cell populations (lower), including Enterocyte (dark green), Enterocyte progenitor (light green), Goblet (dark purple), Immature Goblet (light purple), Tuft (yellow), Stem/TA (dark blue), and Stem (light blue). **B** Diffusion map with additional histograms of first and second dimensions/axes coloured by epithelial cell types. Colour code as in A, with the addition of genomically cancer cells (red). **C** Stacked bar plots of the epithelial cell type compositions across binned diffusion map dimension 2 in normal sample and genomically normal cells, as indicated. **D** UMAP coloured by Cytotrace developmental pseudotime, from early (0, yellow) to late (1, dark purple) in pseudotime space. **E, F** Violin plots of Cytotrace pseudotime across epithelial cell types and consensus call groups, as indicated.

We then wanted to infer cell developmental trajectories. For this, we first embedded epithelial cells from normal and cancer samples into a common diffusion map, thereby emphasizing continuous cell distributions (Fig. 5B). In this embedding, diffusion component (DC) 1 was largely correlated to tuft cell identify, whereas DC2 distributed all other cell types along an apparent differentiation axis, with genomically cancer cells occupying one end. Binning the non-cancer cell types along the DC2 axis (Fig. 5C), we observed that genomically normal stem cells from cancer samples occupied a larger range on the DC2 axis compared to stem cells from normal tissue samples. In contrast, while immature goblet cells and enterocyte progenitors were also more frequent among the cancer-adjacent normal epithelium, they were confined to a similar range on the DC2 diffusion axis compared to normal tissue samples. These results were corroborated by ordering the cell lineages along a pseudo-time axis using CytoTrace ^25^ (Fig. 5D, E). Here, stem cells had a wider distribution in the cancer microenvironment samples, whereas all other cell types were distributed in a fashion comparable to normal tissue. It is of note that the cancer sample-specific stem cell zone extending into the developmental trajectory is composed mainly of cluster 9 stem cells (Fig. 5F), derived from CRCs in the left colon. Together, these analyses suggest that the cancer microenvironment affects differentiation trajectories of normal colonic epithelial cells in their vicinity. The primary difference appears to be stabilization of the stem cell transcriptional state, which in a left-sided CRC microenvironment extends further along the developmental trajectory. In addition, proportions of immature to terminally differentiated cell states are shifted towards the immature cell states in vicinity to CRC.

### The CRC tumour microenvironment is enriched for morphogenetic signal interactions

We next analysed potential paracrine interactions that could underlie the observed differences in cell type frequencies and developmental trajectories between the CRC microenvironment and the normal colon. Our dataset contains a high proportion of immune cells and a lower proportion of stromal cells (Fig. 1B). Specifically, among the 31 663 immune cells, 23 433 were derived from cancer, as were 2 054 of the 2 463 stromal cells. We annotated stromal and immune cell types at a medium granularity using established signatures (Fig. 6A, S6B), in order to strike a balance between accuracy and cluster size. We found that among immune cells, monocytes, macrophages and regulatory T cells were most enriched in the cancer samples, while among stromal cells, fibroblasts were overrepresented in the cancer microenvironment.

**Figure 6.**
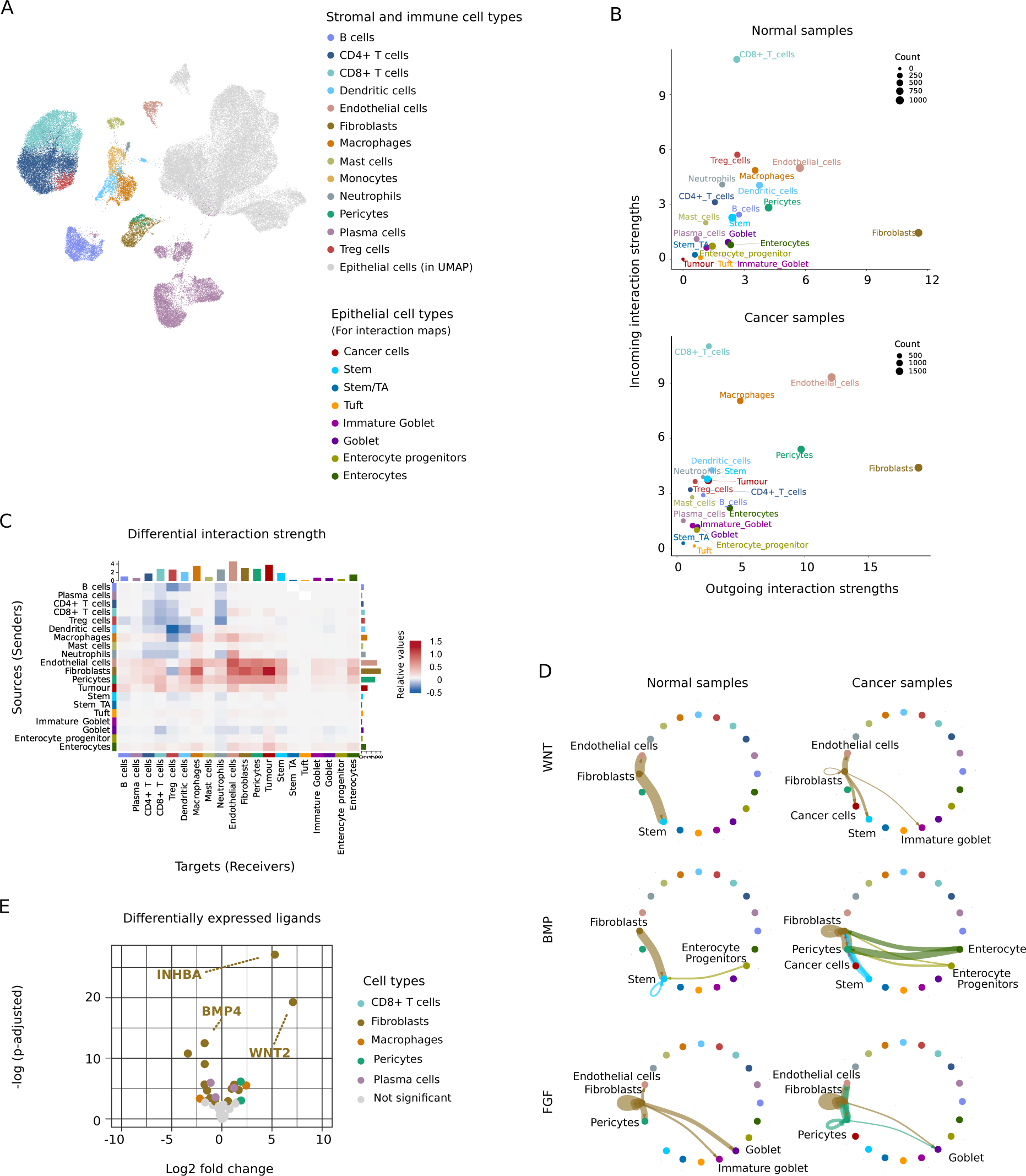
Signalling networks of normal epithelial and genomically normal cells with their respective microenvironments. **A** UMAP of all the cells under analysis, coloured by detailed immune and stromal cell types, and epithelial cells in grey **B-D** Analyses by CellChat **B** Scatterplots of incoming and outgoing signals in normal and cancer samples, as indicated. **C** Heatmap of differential cell-cell communications of cancer samples in contrast to normal samples. **D** Aggregated network graphs of WNT, BMP, and FGF pathways in normal samples versus cancer samples, as indicated. **E** Volcano plot of differentially expressed ligand genes in immune and stromal cell types, as indicated.

We then used CellChat ^26^ to infer interactions in the normal and the cancer samples on a comprehensive basis (Fig. S6C for all interactions). Quantitative analysis revealed that fibroblasts had the most extensive network of outgoing signalling interactions (Fig. 6B) and this network was even larger in cancer samples (Fig. 6C). Endothelial cells and pericytes were rich sources of outgoing signalling interactions in cancer compared to normal. In contrast, endothelial cells, macrophages and pericytes were prominent signal receivers particularly in the cancer microenvironment, whereas CD8+ T cells received the most signals in both, normal and cancer samples (Fig. 6B). Normal epithelial cells emitted and received relatively few signals. Therefore, we analysed key morphogenetic signalling pathway interactions, WNT, BMP and FGF, known to pattern the epithelium in more detail (Fig. 6D). We found that fibroblasts were rich sources of FGF signals potentially received by goblet cells, and of Wnt signals received e.g. by stem cells, and these interactions were seen in both tumour and normal tissue. In addition, BMP interactions known to abrogate the stem cell state ^27^ were diminished in the cancer microenvironment, in particular due to lower BMP expression from fibroblasts. Thus, our data predicts that differences in fibroblast signalling could underlie the changes in normal epithelial cell developmental trajectories that were mainly detected in stem and immature cell population. Indeed, cross-referencing the interactions predicted by CellChat with a curated list of signalling pathway ligands and receptors (Fig. 6E, S6D, S6E; Table S2), we found that WNT2 and the TGF-beta ligand INHBA were most strongly overexpressed by cancer-associated fibroblasts compared to normal fibroblasts, while BMP4 and the WNT co-ligand RSPO3 were expressed at lower levels compared to normal tissue samples.

## Discussion

Single-cell data of cancer tissue often contain transcriptomes of both cancerous and normal epithelial cells. In this study, we used both transcriptome and genome sequence information to trace back the origins of epithelial cell transcriptomes. Across a cohort of CRCs of stages T1-T4 and with different molecular characteristics, a combination of haplotype and SNV level information allowed us to differentiate with high accuracy between cancerous cells and those that are found within cancer tissue but are genomically normal. Using consensus sets of normal and cancer cells, we identified one cluster of genomically normal epithelial cells that was derived from cancer tissue samples exclusively, implying that the cancer microenvironment can result in the adoption of non-standard epithelial cell states in the colon.

Our new tool CCISM makes use of somatic SNVs observed in single-cell sequencing reads for cancer cell identification. While this approach currently requires somatic SNVs independently obtained from matched tumour-normal whole-genome or whole-exome sequencing of the same cancers, it provides the most unambiguous evidence that a cell originated from a cancer lineage. To further benchmark our approach, we also used Numbat, which estimates copy-number variation from shifts in haplotype frequencies over common genetic variants to identify cancer cells, as well as two additional methods that use transcriptome information exclusively. While cancer cell calls from the different approaches show substantial overlap, we find that in our cohort of CRCs the different tools have distinct strengths and limitations contingent on the underlying cancer biology. Therefore, workflows for cancer cell identification should be specifically tailored for the data under analysis. In the final cell annotation of our CRC dataset, cancer and genomically normal cells were largely separated in the underlying louvain cluster structure, implying that cancer and normal epithelial cells do not share common cell states during their developmental trajectories. However, we also observed altered cancer sample-specific cell states in genomically normal epithelial cells, which thus might easily be mistaken for genuine cancer cells. We also caution that our result of largely non-overlapping cell states between normal and cancer may not transfer to cohorts of other stages or types of cancer.

Using the consensus sets of genomically normal and cancer cells defined here, we identified genomically normal *PLA2G2A*-positive stem-like cells arising specifically in the cancer context in the left colon (sigmoid and rectum). PLA2G2A is the human homologue of the gene underlying the mouse Mom-1 locus ^28^, a genetic modifier of familial cancer susceptibility shown to confer cancer resistance in mouse models ^29^. The functional relevance of these stem-like cells remains elusive. On one hand, extension of stem-like and immature cell states along the differentiation trajectory could represent a misguided regenerative process hijacked by paracrine signals of the cancer microenvironment ^30,31^. Indeed, we identify novel paracrine interactions in the CRC microenvironment that were dominated by fibroblasts, as recently also found for breast cancer ^32^. These signals could guide tissue remodelling in the proximity of cancer, which is commonly accompanied by inflammation ^33^. On the other hand, induction of *PLA2G2A*, which we identified as the most specific marker gene of the novel stem-like cells arising near the cancer, could be part of a feedback mechanism to protect the organ from cancer. In agreement with such a function, PLA2G2A is a secreted phospho-lipase that controls tissue homeostasis via modulation of inflammatory responses and is a key player in reducing cancer susceptibility ^34^. The exclusive occurrence of the cancer-induced *PLA2G2A*-positive cells in the left colon suggests regional specificity of the underlying mechanisms along the longitudinal axis of the colon. Supporting region-specific models of cell differentiation, different cell compositions and interaction have been identified in the left-sided/sigmoid colon, such as increased plasma cell interactions ^35^.

Cancer tissue has been shown to extend its influence far beyond its perimeter. Several potential mechanisms with different ranges exist: tumours expressing hormones will affect the complete patient’s body regardless of localization ^36^, while inflammatory responses and other differences in cell composition can have long-range, yet local, effects ^37^. A recent study found prognostic value of gene expression signatures derived from normal-adjacent to CRC issue harvested at a distance of approximately 10 cm from the cancer ^38^, suggesting the existence of long-range interactions between the CRC and surrounding tissues. Thus, gene expression patterns of our normal controls, harvested approximately 10-30 cm from the cancer, may not represent a true normal state, and in extension, our study may underestimate the influence of cancer cells and the cancer microenvironment on adjacent genomically normal colon cells.

New technological developments constantly change single-cell methodology. Employing advances in sequencing depth and transcriptome coverage, e.g., by long read sequencing or specific protocols^39^, a more comprehensive readout of somatic SNVs could be achieved. This would help improve cell lineage determination, e.g., for cancers with few genomic aberrations, such as childhood cancers. With increased coverage, robust *de novo* calling of somatic SNVs could even be feasible directly from single-cell data ^40^. In summary, our study provides general rules for distinguishing between cancer and non-cancer single-cell transcriptomes and provides recommendations how to account for the biology and genetic characteristics of CRC. The rules can easily be adapted for cancers of different origins.

## Methods

### Sample collection and data preprocessing

The sample collection and experimental processing of the clinical specimen for single-cell RNA sequencing data has been described before ^17^ and the new data for P35 was collected and processed using the same protocols. In short, tissues were processed using the Miltenyi Human Tumor Dissociation Kit (Miltenyi, #130-095-929) and a Miltenyi gentleMACS Tissue Dissociator (Miltenyi, #130-096-427), using program 37C_h_TDK_1 for 30–45 min. Single-cell libraries were generated using the Chromium Single-Cell 3’Reagent Kits v3 and the Chromium Controller (10× Genomics). Libraries were sequenced on a HiSeq 4000 Sequencer (Illumina) at 200–400 mio. reads per library. Driver mutations were called as described previously ^17^.

Whole genome sequencing (WGS) data was performed using genomic DNA isolated from microdissected material of snap-frozen (-80°C) CRC tissue, adjacent to material used for single-cell sequencing. DNA was isolated using Qiagen Allprep Kits and sequenced on the Illumina NovaSeq 6000 platform using 2 x 150bp reads. 230-360m reads were generated per sample. Reads were mapped using bwa-mem ^43^ version 0.7.17 against release GRCh38 of the human genome with decoys and virus sequences. For single-cell RNA sequencing data, UMIs were quantified using CellRanger 3.0.2 ^44^ with reference transcriptome GRCh38. See Table S3 for sequencing statistics.

### Single-cell data quality control

All analyses on single-cell data were conducted with Python 3.9.10, Scanpy 1.8.0 ^45^, Numpy seed set at 123, R 4.1.2, and Seurat 4.1.1 ^46^, if not specifically mentioned. CellBender v0.2.2 ^47^ was used to remove ambient RNA with default parameters, 5000 expected cells, and FDR rate at 0.01. We used Scrublet ^48^ for doublet removal and chose the score threshold at 0.3 after inspecting the observed and simulated doublet scores distributions of all the samples. The detected doublet rates ranged from 0.7% to 2.9%. For quality control, cells with min_counts < 1 000, min genes < 500, or mitochondrial percentage > 80% were removed, resulting in a total number of 73 294 cells. The count matrix was then normalized and log1p transformed. The top 2 000 highly variable genes (HVGs) were identified with “patient” as the batch key. Principal component analysis (PCA) was conducted, and we calculated a UMAP using 50 neighbours and 20 principal components.

### Somatic variant calling in WGS and genotyping of single-cell RNA-seq

Somatic variants in whole genome sequencing data were called by Mutect2 from GATK version 4.2.0.0 ^49^ using default parameters. The GATK public resources were used for germline variant loci, common biallelic loci were used to estimate possible contamination, and for the panel of normals. CellSNP-lite^50^ 1.2.2 was used to count somatic variants in single-cell RNA sequencing data against WGS filtered.vcf files with parameters --genotype -p 22 --minMAF 0.001 --minCOUNT 1.

### CCISM model and data simulation

Cancer Cell Identification using Expectation Maximization (CCISM) is a tool for the classification of single-cell expression data based on the expectation-maximization method in Cardelino^20^. Given the total number *d_ij_* of (UMI-collapsed) reads covering variant *i* in cell *j* (reference and variant allele), and the number *a_ij_* of UMIs supporting the alternative allele, we evaluate the likelihood *p_T,j_* that cell *j* is a tumor cell using a binomial model:

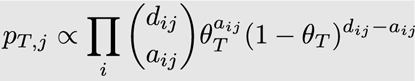

Here, θ*_T_* is the “success probability” for the somatic variants, measuring how likely it is to observe UMIs supporting the variant allele. Similarly, we compute *p_N,j_* as the likelihood that cell *j* is normal, with a fixed nonzero parameter θ*_N_*=0.01 allowing for sequencing errors and uncertainties in the variant calls. We calculate *p_T,j_* and *p_N,j_* in the E-step and estimate the parameter θ*_T_* in the M-step as weighted sum over the counts *d_ij_* and *a_ij_*:

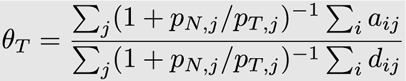

E- and M-steps are iterated until convergence of the likelihood

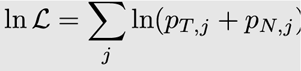

Finally, the likelihoods are normalized to give the posterior cancer cell assignment of a particular cell *p_j_ = p_T,j_ /*(*p_T,j_ + p_N,_*_j_) and a cutoff *p_j_* > .5 is used to define likely cancer cells.

For the benchmark simulations (see also McCarthy et al.^20^), we take the matrix *d_ij_* from a given dataset and simulate values *a_ij_* using a binomial distribution with parameters θ*_T_*=0.4 and θ*_N_*=0.0001 for randomly assigned tumour and normal cell identity, respectively. We used the R package cardelino (v0.6.5) and the BinomMixtureVB function from the vireoSNP package (v0.5.6) for comparison.

### Methodology for consensus cancer calls and trajectory assignments

Epithelial, immune, and stromal cell identity was scored and assigned using previously published cell type markers ^51^. We ran a separate PCA for the epithelial cell compartment and chose 20 neighbours and 15 PCs for the UMAP visualization.

Copy number inference from gene expression profile was performed using inferCNV v1.3.3 ^19^ with default parameters on all the epithelial cells with CellBender-processed ^47^ counts (filtered_h5). The input gene expression profiles were smoothed with a window of 101 genes. The generated dendrograms were cut at k=2 for each patient, and clones were assigned as copy number-aberrant if their averaged smoothed gene expression profile deviated by more than 3 standard deviations from that of clones containing cells of normal samples. Numbat ^22^ 1.0.3 was run with the epithelial cells from the matched normal samples and using default parameters, which included cellSNP-lite 1.2.2 for pile up and Eagle v2.4.1 for phasing the reads. The four samples from P09 (n1, n2, t1, t2) were piled up and phased together, and P26t and P35t were piled-up and phased separately as there were no matched normal samples. The rest of the samples were processed as paired normal and tumour samples.

For iCMS label transfer, we downloaded the CellRanger-processed count matrix (‘Epithelial_Count_matrix.h5 ‘), and the cell-level metadata (‘Epithelial_metadata.csv’) from the source data^11^ (Synapse accession code: syn26844071, https://www.synapse.org/#!Synapse:syn26844071/), filtered by min_genes = 500 and min_counts = 1000, and concatenated this count matrix with ours. The resulting matrix was integrated by scVI with data source as covariate and passed to scANVI to learn the iCMS labels. We found that learning with only the Joanito et al. ^11^ gene list (1 318 genes including a signature for normal cells obtained by personal request from the authors) was suboptimal since it only captured a small proportion of gene expression variance. Therefore, we used the union of all highly variable genes in either dataset and the iCMS signature genes. The resulting matrix was integrated by scVI with data source as covariate and passed to scANVI to learn the iCMS labels.

For the consensus cell identity assignment, we extracted the assignment probability from the outputs of Numbat (p_cnv) and CCISM (CCISM_p), and assigned the cell identity by the following rules: A cell is annotated as genomically cancer cell if (1) p_cnv and CCISM_p are both > 0.5; or (2) CCISM_p = 0.5 and p_cnv > 0.5 ; or (3) p_cnv > 0.5 in MSS samples; or (4) CCISM_p > 0.5 in MSI samples. A cell is annotated as genomically normal cell if p_cnv and CCISM_p are both < 0.5. A cell that does not fit into any of the categories above is annotated as ‘unclear’ and removed from the downstream analysis For detailed epithelial cell type annotation, we used scVI and scANVI to integrate datasets and learn cell type labels from Uhlitz et al. The scVI models were trained on the raw count matrix (adata.layer[‘count’]) of 2000 highly variable genes using scvi-tools 0.19.0 with patient and percent_ribo as covariates. These models were used by scANVI as input to predict cell type labels of newly included cells based on the annotation of previously annotated cells.

The linear mixed model for cell type composition was composed using the ‘glmer’ function with binomial distribution from the lmer package ^52^. For each cell type, we tested if there is a difference between genomically normal cells and healthy cells from normal samples, where patient was included as a random effect variable.

To enhance concrete transcriptomic contrasts between cancer and normal cells, 1498 cells from normal samples but were assigned as tumour-centric cell type, namely TC1-4, were removed from the downstream analysis. The epithelial cell type of genomically cancer cells was then assigned as ‘cancer-like’ in transcriptomic analysis. Diffusion maps were calculated with 15 neighbours and CytoTrace pseudotime as implemented in CellRank 1.5.2.dev236+gab03900 ^53^.

### Methodology for scoring CRC signalling pathways and inferring paracrine interactions

We curated a list of known ligands and receptors of key signalling pathways in CRC and a list of CRC signature genes for specific phenotypes from literature (Table S1). The expression levels of CRC signatures were calculated using ‘score_gene’ function in Scanpy. The paracrine interactions within normal and tumour samples were inferred by CellChat 1.6.1.

### Data and Code availability

Processed single-cell RNA sequencing data and somatic variant allele counts are available on zenodo via doi:10.5281/zenodo.10692019. CCISM is available from github.com/bihealth/CCISM. Analysis code is available from github.com/bihealth/Wei_et_al_2024.

### Ethics Permission

All patients were aware of the planned research and agreed to the use of tissue. Research was approved by vote EA4/164/19 of the ethics commission of Charité—Universitätsmedizin Berlin.

### Author contributions

D.B., M.M. and B.O. designed research; D.H., C.S., N.B., D.B., M.M. and B.O. supervised research; T.-T.W., E.B., S.P., M.M. and B.O. analyzed data; P.B. and A.T. performed experiments; M.M. and B.O. wrote the manuscript with input from all other authors. The work reported in the paper has been performed by the authors, unless clearly specified in the text. The authors have no conflict of interests related to this publication.

## Acknowledgements

We thank Edda von der Wall and Hedwig Lammert (Charité, Institute of Pathology) for excellent technical assistance. The work was in part funded by Deutsche Forschungsgemeinschaft (RTG CompCancer GRK2424/1) and by the BIH-funded PeDiOn and Clinical Scientist programs. We acknowledge excellent services by the BIH Sequencing core facility.

