## Supplementary Figures for "High-confidence calling of normal epithelial cells allows identification of a novel stem-like cell state in the colorectal cancer microenvironment"

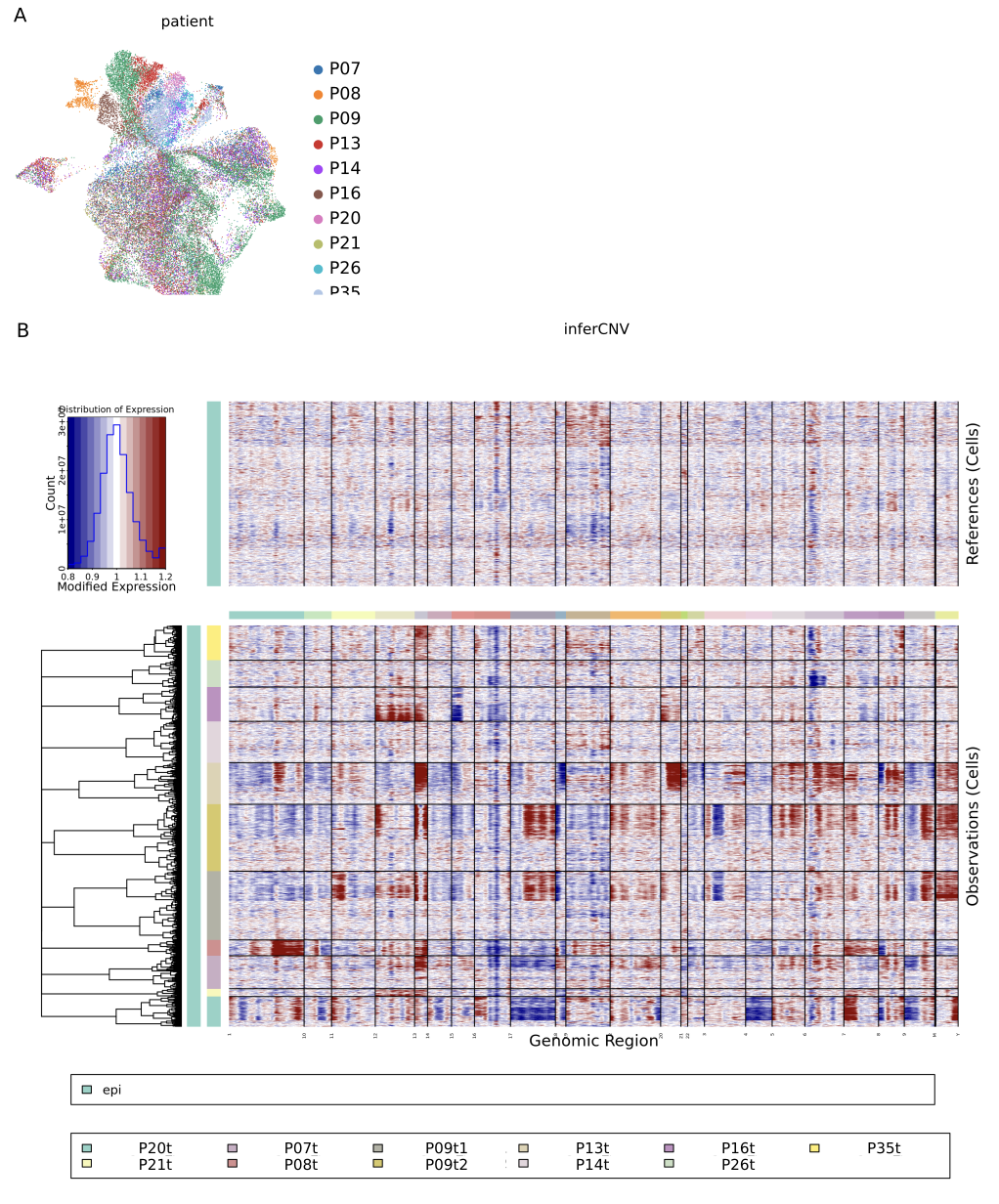

**Supplementary Figure 1**: **A** UMAP plot of epithelial cells coloured by patient. **B** Heatmap of inferCNV result: upper panel shows the modified expression level of normal epithelial cells as reference and lower panel shows the modified expression level of epithelial cells from tumour samples

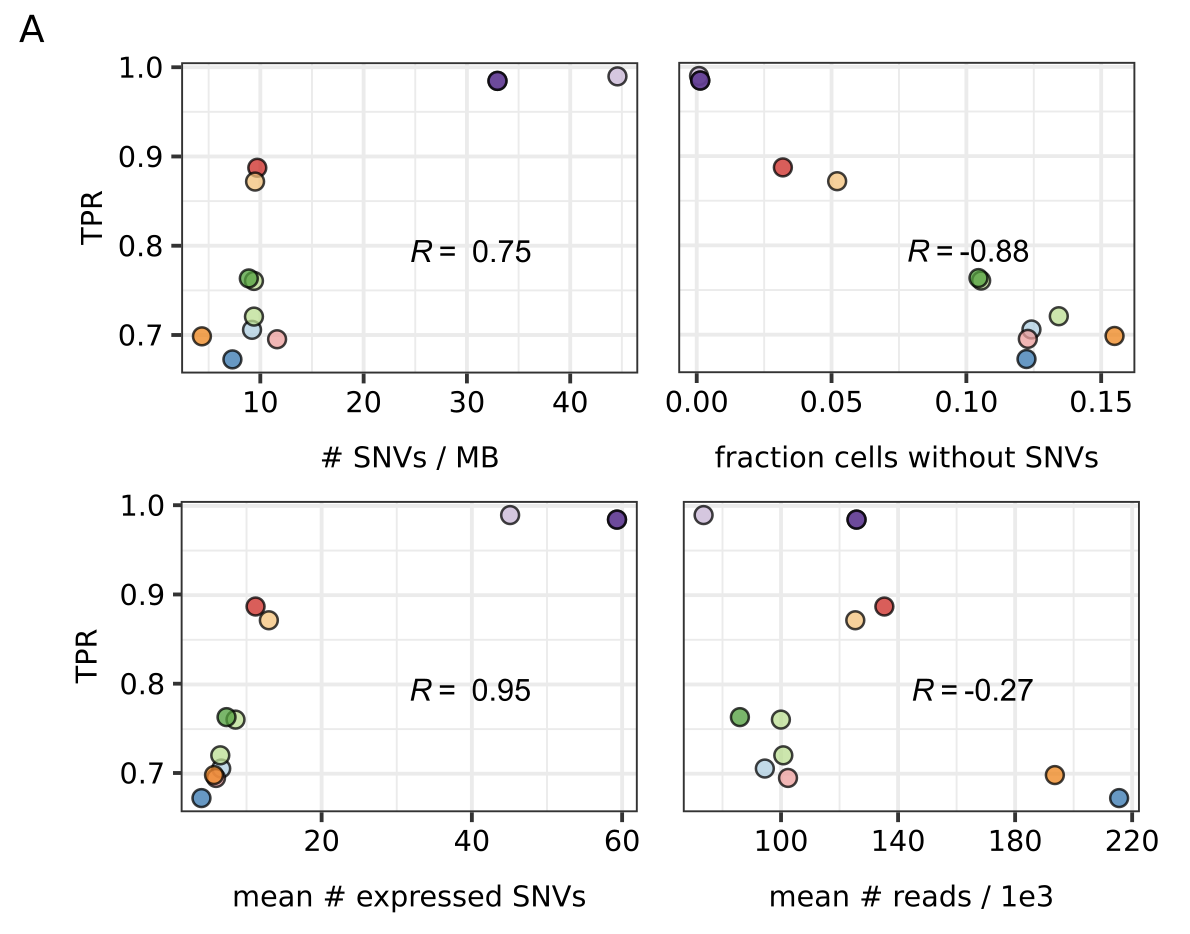

**Supplementary Figure 2**. Scatterplots of the true positive rates (TPR) of CCISM in four conditions across samples: number of SNV per MB (upper left), fraction of cells without SNVs (upper right), average number of expressed SNVs (lower left), and average number of read scaled by 1000 (lower right)

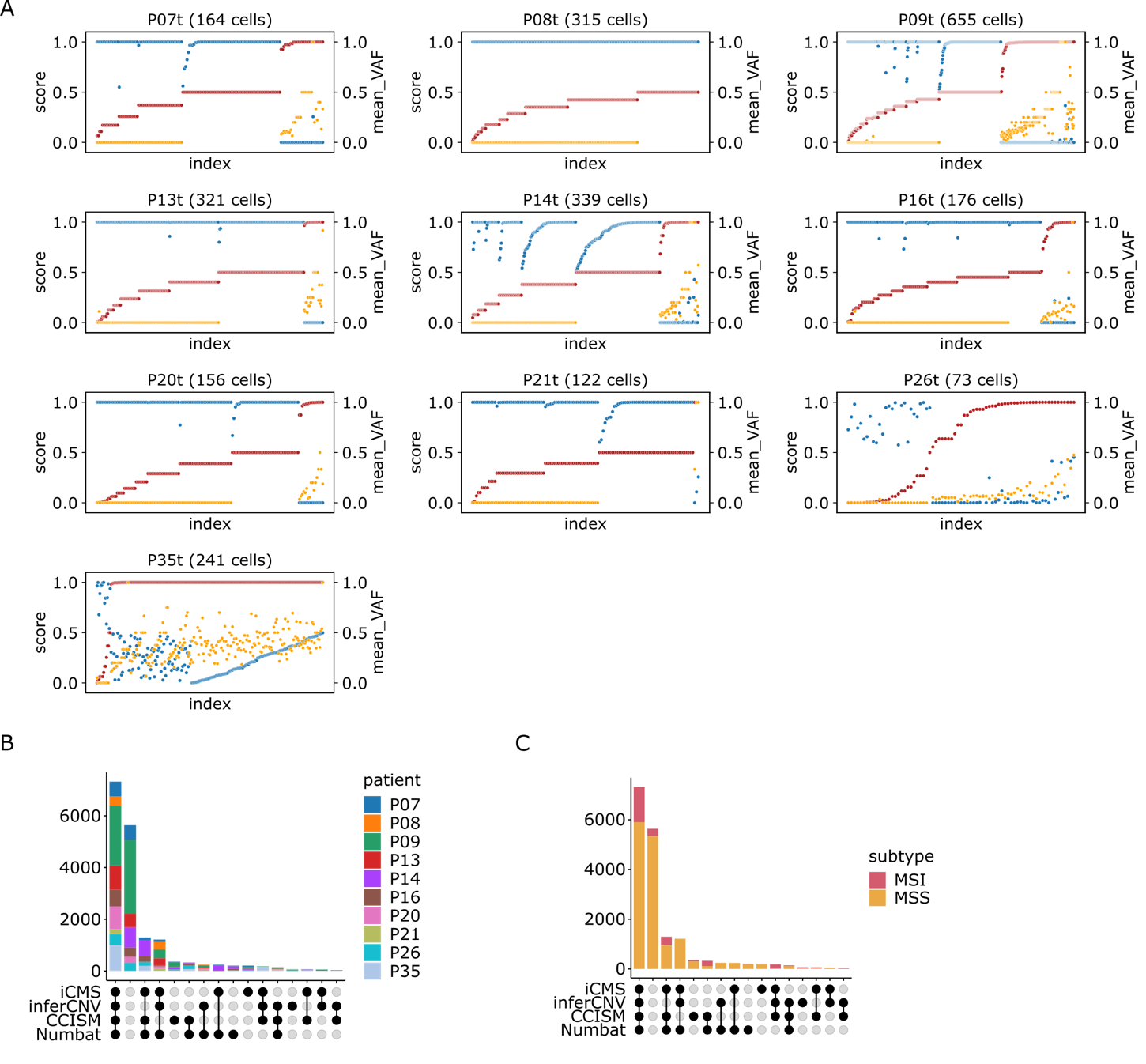

**Supplementary Figure 3. A** Scatterplots of cells with disagreements in CCISM score (red, left y axis) and Numbat score (blue, left y axis), with mean variant allele frequency (orange, right y axis) per cell across tumour samples. Each plot was ordered by the ascending CCISM score, then the ascending Numbat score. **B** Upset plot of the intersection of cancer cell calls between iCMS, inferCNV, CCISM, and Numbat coloured by patient (colours match with Fig1H). **C** Upset plot of the intersection of cancer cell calls between iCMS, inferCNV, CCISM, and Numbat coloured by microsatellite status (MSI in red and MSS in yellow).

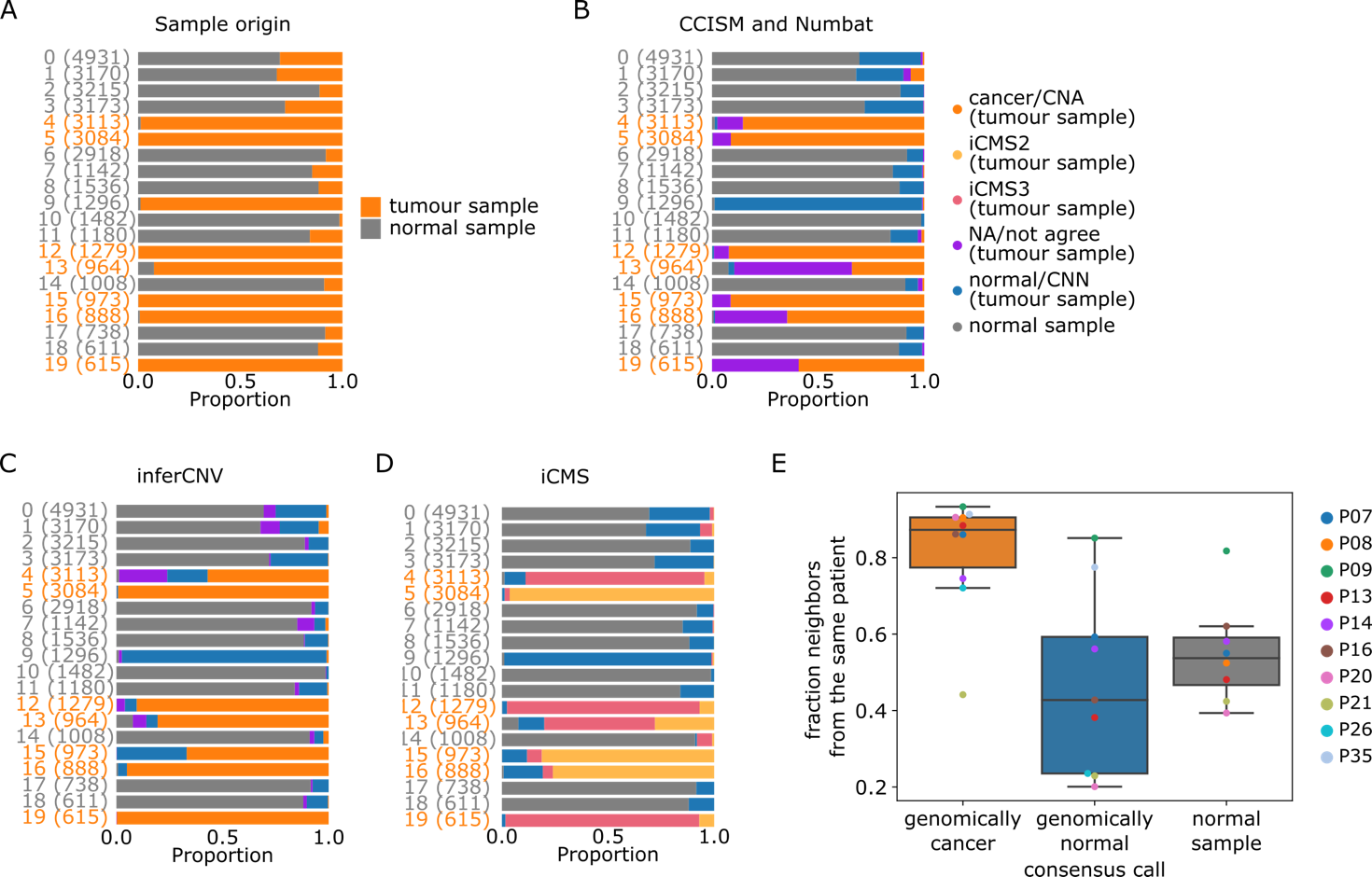

**Supplementary Figure 4. A** Bar plot of sample origin distribution of each cell across each louvain cluster (tumour sample in orange and normal sample in grey). **B** Bar plot of the intersection of CCISM and Numbat cancer cell calls across louvain cluster. Cancer cells were coloured as orange, cells which CCISM and Numbat do not agree as purple, normal cells as blue, and cells from normal samples as grey. **C** Bar plot of copy number status calls by inferCNV (CNA in orange, NA in purple, CNN in blue, and cells from normal samples in grey). **D** Bar plot of the cancer cell calls by iCMS where iCMS2 calls were in yellow, iCMS3 calls in pink, normal calls in blue, and cells from normal sample in grey. **E** Boxplot of the mean fraction of neighbours from the same patient per cell, coloured by patient (colours matched with Fig1H)

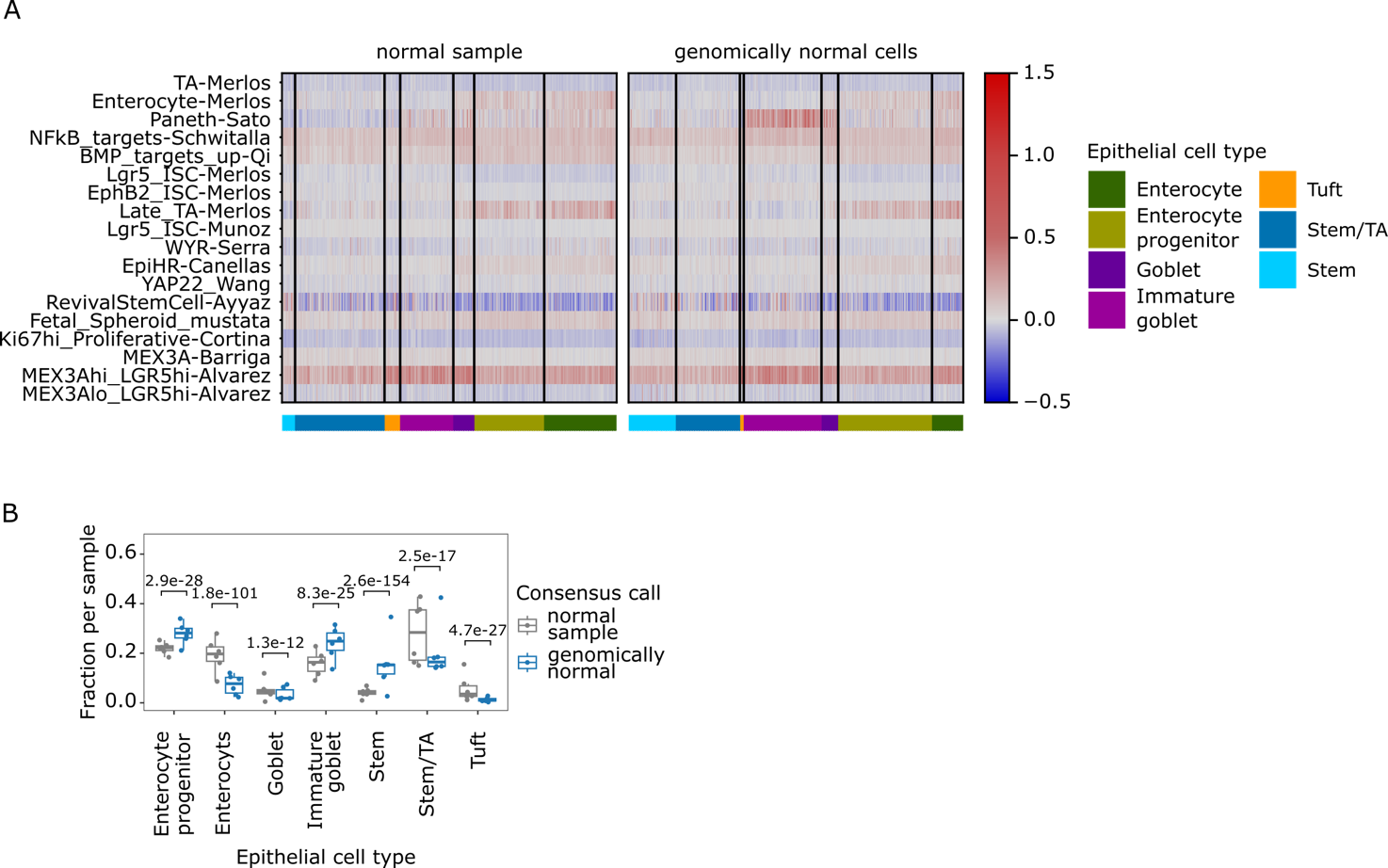
**Supplementary Figure 5. A** Heatmaps of the average gene expression for CRC signature pathways in a curated list from the literature, normal samples on the left, genomically normal cells on the right. **B** Boxplot of cell type fractions in normal samples vs. genomically normal cells. P-values from mixed-effects binomial model.

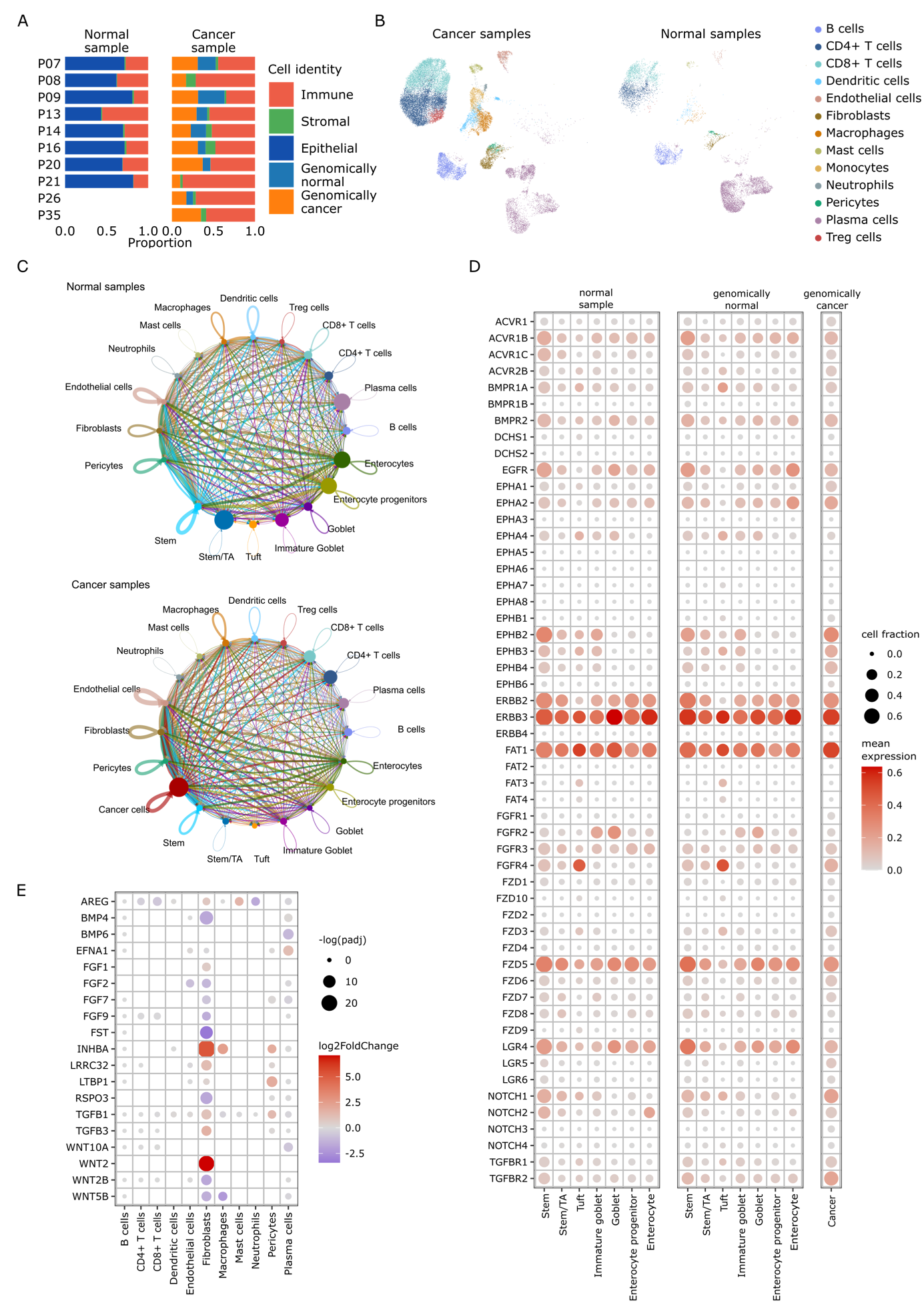

**Supplementary Figure 6. A** Bar plots of cell class distributions across patient and sample origin, including immune cells in dark orange, stromal cells in green, epithelial cells from normal sample in dark blue, genomically normal cells from tumour sample in teal, and genomically cancer cells in light orange. **B** UMAPs including immune and stromal cells from tumour (upper) and normal samples (lower). **C** Aggregated network graphs of inferred cell-cell communication strength by CellChat in normal and cancer samples, as indicated **D** Dot plots of mean expression levels (colour) of receptor genes and fractions of cell expressed the receptor genes (size) across normal sample, genomically normal cells, and genomically cancer cells. **E** Dot plots of differentially expressed ligand genes in immune and stromal cells (colour: log2 Fold Change of genomically normal cells vs. cells from normal samples, size: adjusted p values from a differential gene expression test by DESeq2).
